# Decoding 3D spatial location across saccades in human visual cortex

**DOI:** 10.1101/2020.07.05.188458

**Authors:** Xiaoli Zhang, Christopher M Jones, Julie D Golomb

## Abstract

Visual signals are initially processed as two-dimensional images on our retina, but we live in a 3D world. Depth information needs to be reconstructed from the 2D retinal images, using cues such as binocular disparity. But in daily life, we also make frequent, rapid eye movements, which alter the 2D retinal input. How do we achieve stable 3D perception across saccades? Using fMRI pattern analysis, we investigated how 3D spatial representations in human visual cortex are influenced by saccades. Participants viewed stimuli in four possible 3D locations, defined by 2D vertical position (above or below screen center) and depth position (in front of or behind central screen plane). We compared the amount of 2D and depth information in visual cortical regions during no-saccade blocks (stationary fixation) with that during saccade blocks (series of guided saccades). On no-saccade blocks, decoding of stimulus location was highly *dependent* on fixation position: in later visual areas we could decode both vertical and depth information across blocks that shared the same fixation position (as previously reported), but little vertical or depth information could be decoded across blocks with *different* fixation positions. Strikingly, the neural similarity patterns appeared *tolerant* to changes in fixation position during saccade blocks: despite the saccade-induced retinal and fixation changes, we could reliably decode both vertical and depth information. The findings suggest that representations of 3D spatial locations may become more tolerant of fixation positions during dynamic saccades, perhaps due to active remapping which may encourage more stable representations of the world.

**Significance:** This study investigates two fundamental challenges for visual perception: how to preserve spatial information across frequent eye movements, and how to integrate binocular depth location with 2D location to form coherent 3D percepts. Aspects of these challenges have been studied in isolation, but surprisingly no studies have investigated them jointly to ask how 3D spatial representations in human visual cortex are influenced by saccades. Our fMRI pattern analysis findings highlight a potentially critical role of active, dynamic saccades on stabilizing 3D spatial representations in the brain, revealing that representations of 3D space may be modulated by eye position during sustained fixation, but could become tolerant of changes in eye position during active, dynamic saccades.

## Introduction

Visual inputs are initially processed on the retina in two-dimensional, eye-centered coordinates, which provides direct coding of 2D spatial locations from the very early stages of visual processing. But our perceptual experience is of a 3D world, and one that is stable across eye movements. We move our eyes around frequently, on the order of a few times per second. As a result, the retinal positions of objects projected to our eyes also change frequently, creating great challenges for visual cognition, and an extra layer of complexity for 3D perception.

Many studies have explored different aspects of this challenge. For example, 2D perception across saccades is known to be disrupted in multiple ways. Objects flashed around the time of a saccade tend to be systematically misperceived as closer to the saccade endpoint than they actually are, a phenomenon called compression of space (1–3). Localization of a foveal target is also worse when performed after a saccade compared to without a saccade, and responses tend to be biased towards the former fixation location (4). Furthermore, saccades can interfere with spatial attention (5, 6), memory (7–9), and feature perception, including an effect where the features of objects at two different locations can be mixed (10, 11). An important question in vision research has been how our brain compensates for the disturbance from executing saccades and maintains stability, via remapping and other mechanisms (see reviews 12–14).

Other studies have explored the mechanisms of depth perception. Our retinal images are only two-dimensional, which means that in order to perceive accurately the three-dimensional world, we need to reconstruct the third dimension, depth, from the 2D retinal inputs. Depth can be reconstructed from many visual cues, such as size, perspective, shading, motion parallax, and binocular disparity (15). Among these cues, binocular disparity (small horizontal differences in an object’s projected location on the two eyes) is particularly effective (16). Instead of perceiving two images of the same stimulus, our brain is able to fuse the two images and perceive how far the stimulus is based on how separated the retinal images are (i.e., stereoscopic vision).

In terms of neural representations, it is widely shown that 2D spatial information is represented throughout visual cortex and beyond (17–23). More controversial are the reference frames of 2D spatial representations across saccades. Some studies have shown that these representations are primarily coded in retinotopic (eye-centered) coordinates, with no evidence for explicit spatiotopic (world-centered) representations (24–26). However, other studies have shown some evidence for spatiotopic updating or adaptation across saccades (27–31).

Studies examining the neural representations of depth perception have typically focused on sensitivity to binocular disparity (32–36) and other depth cues (37–40) (also see review 41). A few recent studies have attempted to explore the nature of *3D* spatial representations in the human brain by varying both 2D location and position in depth: A study from our lab revealed that multivoxel pattern information about position-in-depth increases along the visual hierarchy while information about 2D location (horizontal and vertical locations) decreases; that is, representations of spatial locations transition from 2D-dominant in early visual areas to balanced 3D in later visual areas (33). Another study using inverted encoding models similarly recovered representations of both 2D and depth position in visual and parietal areas, particularly in V3A (34).

However, no studies have investigated how these 3D spatial neural representations are affected by eye movements. Behaviorally, there is some evidence that executing saccades creates challenges for depth processing, which may not be surprising given that reconstructing depth from binocular disparity relies on precise retinal position information, and retinal position changes with each eye movement. Horizontal eye movements in particular can interfere with our processing of depth, e.g., biasing the depth perception of stimuli which flash around the time of a saccade to be closer (42), similar to 2D mislocalization, and impairing memory-guided reaching in depth (43). Interestingly, however, there is also evidence that self-generated motor actions, including eye movements, can *enhance* our perception of 3D space (44).

In the current study, we examine how 3D spatial representations in human visual cortex are influenced by saccades. We modified the fMRI paradigm from Finlayson et al. (33), where participants wore red-green anaglyph glasses in the scanner and viewed a random dot stimulus that appeared in one of four 3D locations (Figure 1), varying in 2D position (above or below screen center; vertical information), and depth position (in front of or behind central screen plane). Participants either remained fixated at a single fixation location (left or right of screen center) for 16 sec (no-saccade blocks) or made a sequence of back and forth saccades (initially starting from left or right of screen center) over the 16 sec (saccade blocks); they were instructed to passively view the 3D stimulus in the periphery. We used multivariate pattern analysis (MVPA) to investigate the representations of 2D (vertical; y-axis) location information and depth (z-axis) location information in different visual regions and compared the representations of 3D location during sustained fixation and dynamic saccade blocks.

**Figure 1.**
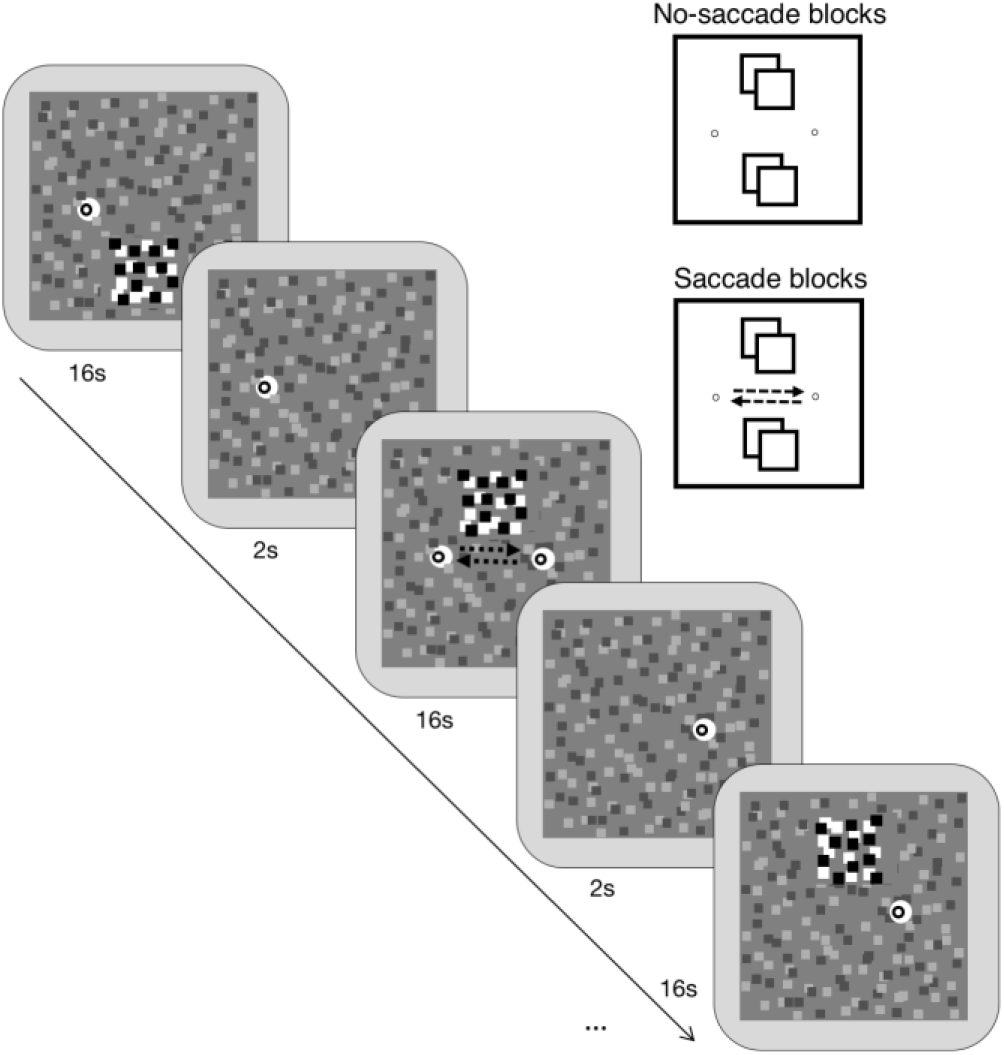
Experiment paradigm. Each block lasted 16s, with 2s inter-block interval. In each block, a dynamic RDS patch stimulated one of four locations, defined by its vertical location (above or below the screen center) and its depth location (in front of or behind the screen depth plane). In half of the blocks, participants kept fixated at the fixation dot at either left or right of the screen center throughout the block; in the other half, participants made repetitive saccades between the left and right, following the alternating fixation dot (as shown in the schematics in the upper right).

## Results

### Vertical and depth location information in visual regions of interest (ROIs)

Similar to Finlayson et al.(33), we used an fMRI pattern similarity split-half correlation approach to calculate the amount of MVPA information about a stimulus’ vertical and depth location, within each pre-defined ROI group (early visual areas V1, V2, and V3; intermediate visual areas V3A, V3B, and V4; later visual areas V7, V8, and IPS; and category selective areas LOC and MT+), for each type of analysis (details in Materials and Methods). To quantify the main effects and interactions, we submitted the data to 2 (location type: Y and Z) × 4 (ROI group) repeated measure ANOVAs, and then we performed post-hoc comparisons quantifying the amount of Y and Z information present in different ROIs. Below we report the data for the grouped-ROIs; individual ROIs and searchlight data are presented in the supplemental materials.

As an initial analysis, we collapsed across both saccade and no-saccade blocks, as well as left and right fixation positions in no-saccade blocks and initial fixation positions in saccade blocks (Figure 2) to get a sense of overall 3D (vertical and depth) information. There was a significant main effect of location type, *F*_1,11_=35.51, *p*<.001, *η_p_^2^*=.764, indicating greater information about vertical than depth locations across all visual ROIs. Furthermore, we found a significant main effect of ROI group (*F*_1.416, 15.580_=15.71, *p*<.001, *η_p_^2^*=.588), and a significant interaction between location type and ROI group (*F*_1.299,14.286_= 18.89, *p*<.001, *η_p_^2^*=.632). Post-hoc one-way ANOVAs showed that vertical location information decreased along the visual hierarchy, *F*_1.329,14.621_=17.76, *p*<.001, *η_p_^2^*=.618, whereas depth location information increased, *F*_1.819,20.008_=5.382, *p*=.015, *η_p_^2^*=.329. Post-hoc t-tests showed that both vertical and depth information were significant in later visual (V7/V8/IPS) and category-selective (LOC/MT+) groups of ROIs, *t*’s≥2.804, *p*’s≤.017, Cohen’s *d*’s≥0.809. These results replicate the findings in Finlayson et al.(33).

**Figure 2.**
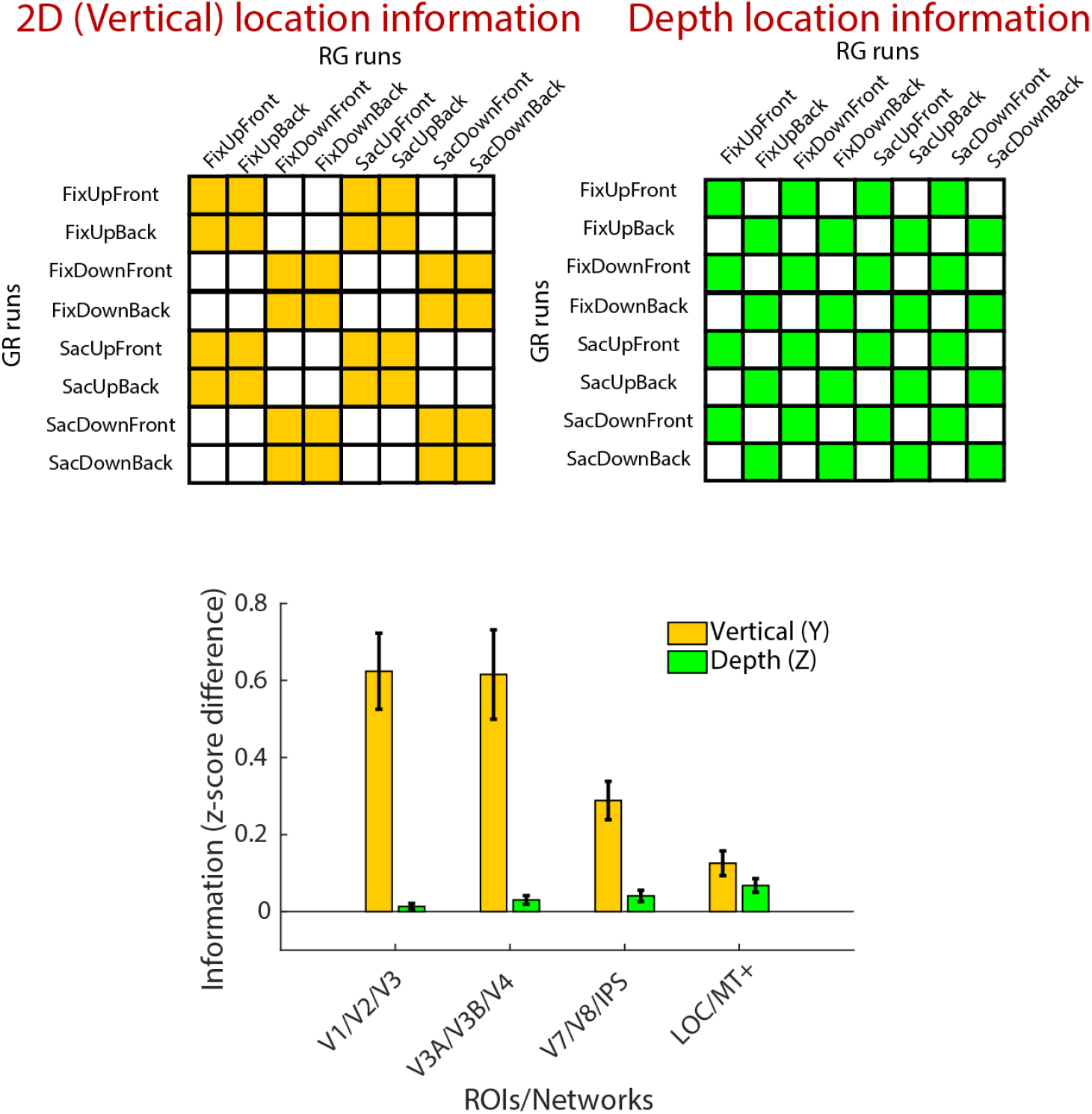
3D location information from MVPA with data from all blocks. Top row shows correlation matrices used for calculating overall Y/vertical (left panel) and Z/depth (right panel) location information, using data from all blocks. Location information is calculated by subtracting between-category correlation coefficients (white cells) from within-category coefficients (colored cells). Bottom row shows the results, with Y and Z information for the four ROI groups. Error bars represent between-subject *SEM*.

Our primary research question was how 3D location information is represented in dynamic saccade blocks, compared to in no-saccade blocks. We thus separately calculated location information in no-saccade and saccade blocks (Figure 3A & 3B), and performed a 2 (saccade condition: saccade and no-saccade) × 2 (location type: Y and Z) × 4 (ROI group) three-way repeated measures ANOVA. We again found a significant main effect of location type (*F*_1,11_=33.353, *p*<.001, *η_p_^2^*=.752), a significant main effect of ROI group (*F*_1.528,16.804_=15.661, *p*<.001, *η_p_^2^*=.587), and a significant interaction between location type and ROI group (*F*_1.326,14.584_=17.645, *p*<.001, *η_p_^2^*=.616). Most importantly, there was no significant main effect of saccade condition in the three-way ANOVA, *F*_1,11_=1.046, *p*=.328, *η_p_^2^*=.087, nor were any interactions with saccade condition significant (all *F*s≤1.394, *p*s≥.270, *η_p_^2^*s≤.112). Post-hoc t-tests confirmed that in saccade blocks, both vertical and depth information were significant in later areas and category-selective areas (*t*’s≥2.417, *p*’s≤.034, Cohen’s *d*’s≥0.698). This result revealed that both vertical (2D) and depth information in these later areas was preserved in saccade blocks, such that making dynamic saccades did not seem to impair the 3D location information at all, compared to in no-saccade blocks.

**Figure 3.**
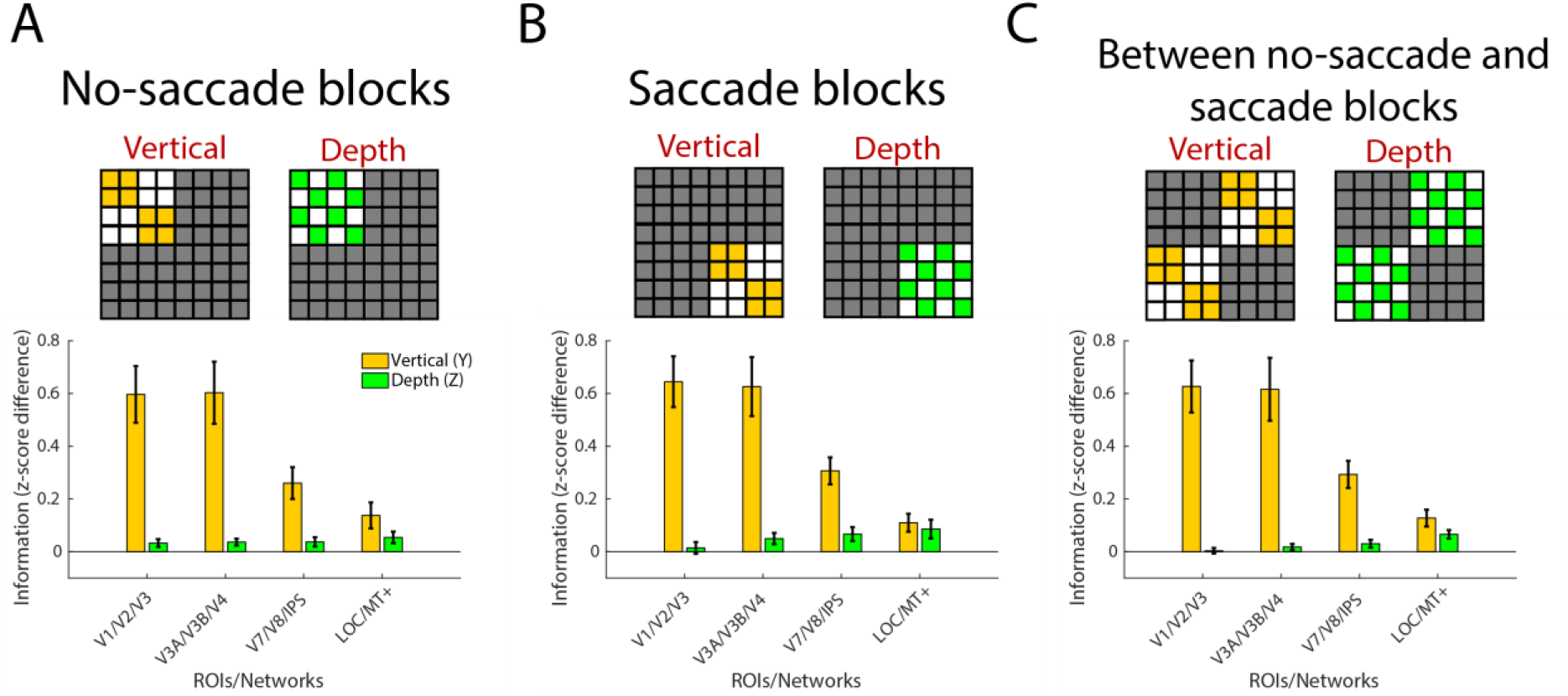
3D location information, separating no-saccade and saccade blocks. Y and Z information are calculated in the same way as depicted in Figure 2 (within-category minus between category), but here only from subsets of the cells in the correlation matrices (unused cells shown as gray). (A) Location information calculated with data from no-saccade blocks only. (B) Location information calculated with data from saccade blocks only. (C) Cross-decoding location information calculated using MVPA correlations between no-saccade and saccade blocks. Error bars represent between-subject *SEM*.

We further asked whether similar brain patterns for location were shared between no-saccade and saccade blocks, or if distinct brain patterns were used to represent spatial location separately during fixation and across saccades. To answer this question, we attempted to cross-decode location information in the two contexts, by correlating the brain patterns *between* no-saccade and saccade blocks. As shown in Figure 3C, the cross-block MVPA results were similar to the results within no-saccade blocks or saccade blocks alone: There was a significant main effect of location type (*F*_1,11_=34.75, *p*<.001, *η_p_^2^*=.760), a significant main effect of ROI group (*F*_1.361,14.975_=15.06, *p*<.001, *η_p_^2^*=.578), and a significant interaction (*F*_1.3611, 14.416_=19.35, *p*<.001, *η_p_^2^*=.638). Vertical location information could be significantly cross-decoded in all four ROI groups (*t*’s≥3.997, *p*’s≤.002, Cohen’s *d*’s≥1.154), and depth location information could be significantly cross-decoded in LOC/MT+ (*t*_11_=4.129, *p*=.002, Cohen’s *d*=1.192).

### Tolerance of location information across fixation positions and saccade directions

The above analyses collapsed across fixation blocks with both left and right fixation positions and saccade blocks with different initial fixation positions (which also mean different saccade directions; see Materials and Methods for details). This procedure could have contributed to the decoding and cross-decoding results above. An important question is then whether the observed vertical and depth location information is *dependent* on fixation position and/or saccade direction, or whether it is tolerant of these differences. In a second stage of analyses, we broke down these conditions to compare within-versus across-fixation location information in no-saccade blocks, and within-versus across-saccade-direction location information in saccade blocks (Figure 4).

**Figure 4.**
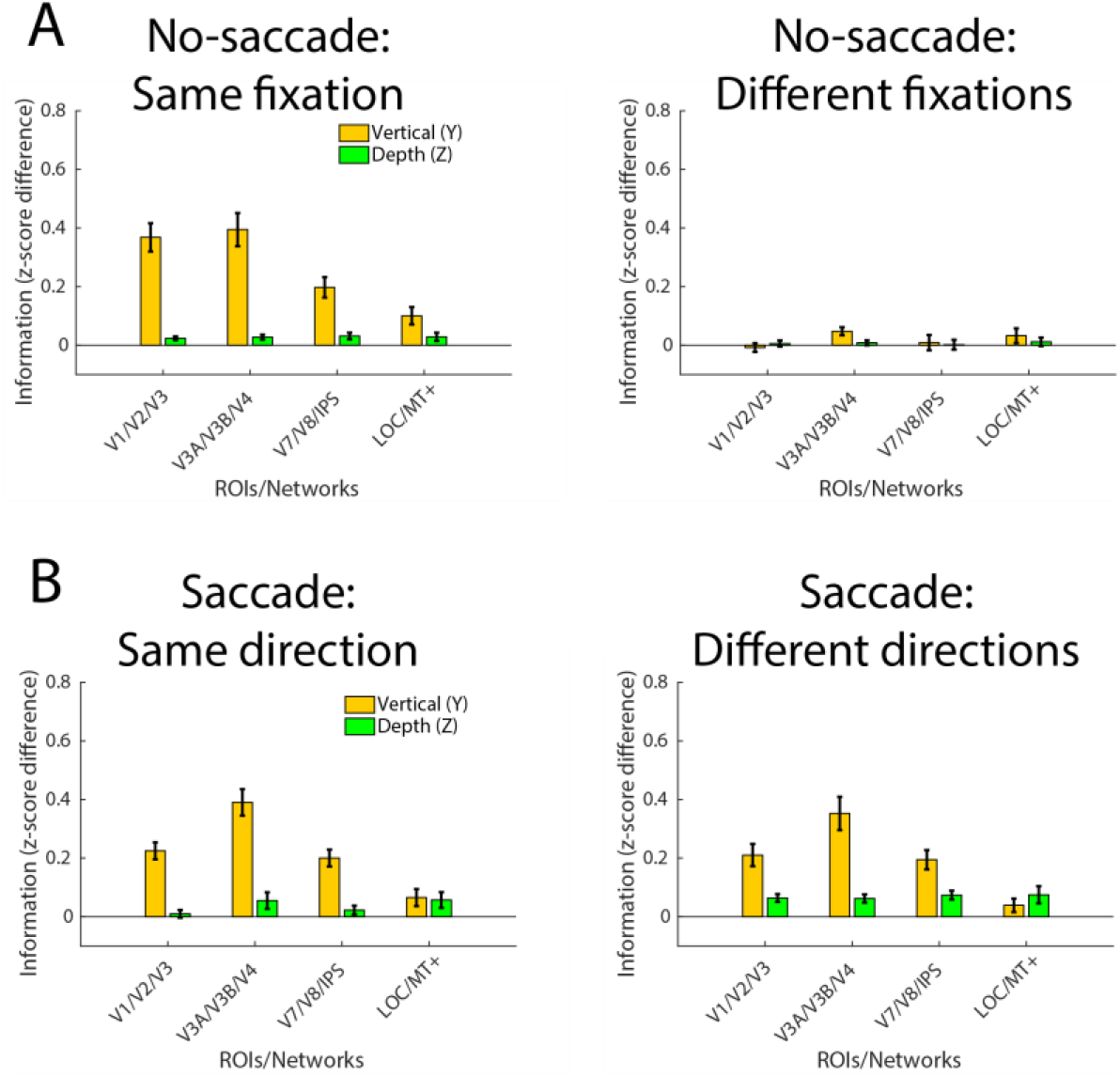
Tolerance/dependence analysis: Vertical (Y) and depth (Z) location information calculated separately for same vs different fixation position (no-saccade blocks) and saccade direction (saccade blocks). (A) No-saccade blocks. Y and Z location information calculated from MVPA between blocks with the same fixation position (left panel), and different fixation position (right panel). Y and Z location information is dependent on fixation position in no-saccade blocks. (B) Saccade blocks. Y and Z location information calculated from MVPA between saccade blocks with the same saccade directions (left panel), and different saccade directions (right panel). Y and Z location information is tolerant of fixation pattern differences in saccade blocks. Error bars represent between-subject *SEM*.

To test if the representations of vertical and depth location are tolerant of fixation position during periods of stable fixation, we separated the no-saccade blocks with left and right fixation positions, and then analyzed vertical and depth information within blocks that shared the same fixation position (“same-fixation”), and separately between blocks of different fixation positions (“different-fixation”). As shown in Figure 4A, we found significant MVPA information about both vertical and depth location in no-saccade blocks sharing the same fixation position, but not across blocks with different fixation positions. We ran a 2 (fixation position: same-fixation and different-fixation) × 2 (location type: Y and Z) × 4 (ROI group) three-way repeated measures ANOVA. In addition to the significant main effects of location type and ROI group, there was also a significant main effect of fixation position, as well as significant interactions between fixation position and location type, fixation position and ROI group, and among all three variables (all *F*s≥10.97, *p*s<.001, *η_p_^2^*s≥.499). This suggests that the representations of both vertical and depth location were largely dependent on fixation position in no-saccade blocks.

On the other hand, when we separated the saccade blocks with left and right initial fixation positions (i.e., different saccade directions), we found a different pattern (Figure 4B). We again found the standard significant main effects of location type and ROI group, as well as a significant interaction between location type and ROI group (*F*s≥18.922, *p*s<.001, *η_p_^2^*s≥.632). However, here there was no significant main effect of saccade direction (*F*_1,11_=0.072, *p*=.793, *η_p_^2^*=.007); information about both vertical and depth location were comparably maintained during saccade blocks that shared the same saccade direction pattern and across blocks with different saccade directions (i.e. different fixation locations at each point in the block). There was a significant interaction between saccade direction and location type, *F*_1,11_=22.277, *p*<.001, *η_p_^2^*=.669, but all other interactions were not significant, *F*s≤1.816, *p*s≥.197, *η_p_^2^*s≤.142.

The post-hoc t-tests revealed that vertical and depth location information were significant for most ROI groups and comparisons, but the individual t-tests were less reliable^1^, and the magnitudes of both vertical and depth information were smaller than in the previous collapsed analyses. A potential reason is that this analysis was underpowered compared to the initial set of analyses due to looking at only a subset of the whole dataset (i.e., 1/8 of the power compared to Figure 2), and the single-run GLMs required for the saccade direction breakdown may have additionally introduced more noise, compared to the analogous no-saccade breakdown (details in Materials and Methods). However, while lower power might be expected to make it harder to detect depth location information (which was weaker to begin with), the vertical location information is quite robust, so the near-total lack of vertical location information in the different-fixation condition is striking (i.e. comparing the yellow bars in the right panel of Fig 4A to the other three panels in Figure 4). Indeed, a post-hoc omnibus ANOVA on the y-information revealed a significant saccade presence (saccade vs no-saccade) × tolerance (same vs different. fixation/saccade direction) interaction, with a robust effect size, *F*_1,11_=19.568, *p*=.001, *η_p_^2^*=.640. This suggests that information about the stimulus’ location was tolerant of saccade direction and changes in fixation position in the dynamic saccade blocks, but was not tolerant of fixation position in the static no-saccade blocks.

### Timepoint-by-timepoint tolerance analysis

The above analyses revealed that vertical and depth location information could be decoded during saccade blocks, and appeared largely tolerant of saccade direction. This is particularly interesting in comparison to the lack of tolerance to fixation position found in the no-saccade blocks. The different saccade directions by definition involve different fixation locations; in other words, saccade blocks could be considered as a series of alternating short periods (2 s) with left and right fixation positions. When comparing the two different saccade directions, at any given point in the block, the fixation position is different for a sacLR versus sacRL block. The fact that stimulus location information was tolerant to changes in fixation position in the saccade blocks but not the no-saccade blocks is a striking result. One explanation we entertain later in the discussion is that this difference could be due to the dynamic, active context of the saccade blocks; i.e. that the saccades trigger a more tolerant representation. But there are alternative explanations that could stem from artifacts of the block design: The GLMs used in the above analyses modeled the whole 16 second block as a single event, which could have effectively blended or combined activity across the temporal sequence of left and right fixations. In other words, it’s possible that the apparent tolerance across saccade direction could have been an artifact of the activation patterns containing location information from *both* left and right fixation positions during saccade blocks, regardless of saccade direction.

To address this possibility, we performed an MVPA time-course analysis, examining the MVPA contrasts reported above (comparing vertical and depth information for same fixation location, different fixation location, same saccade direction, and different saccade direction), but now on activity patterns for each timepoint, estimated from FIR GLMs (see Materials and Methods). Because this is a lower powered analysis, it may be ill-suited to detect small effects, and is presented primarily as an exploratory analysis. Indeed, the magnitude of depth (z) information was too small to draw reliable conclusions in this timepoint-by-timepoint analysis (supplemental Figure S3), so we focus here on the larger vertical (y) information effects. MVPA timecourses are shown in Figure 5: in the early, intermediate, and later visual areas (those regions in which we could best decode vertical location information in the whole-trial MVPA), the timecourse of decoding peaks after several seconds and remains sustained for the block, for three out of the four contrasts. Critically, vertical information during saccade blocks could be successfully decoded on a timepoint by timepoint basis even across blocks with different saccade directions (different fixation positions at each timepoint). Conversely, vertical information was not tolerant of different fixation positions during no-saccade blocks, consistent with the whole-trial results reported above.

**Figure 5.**
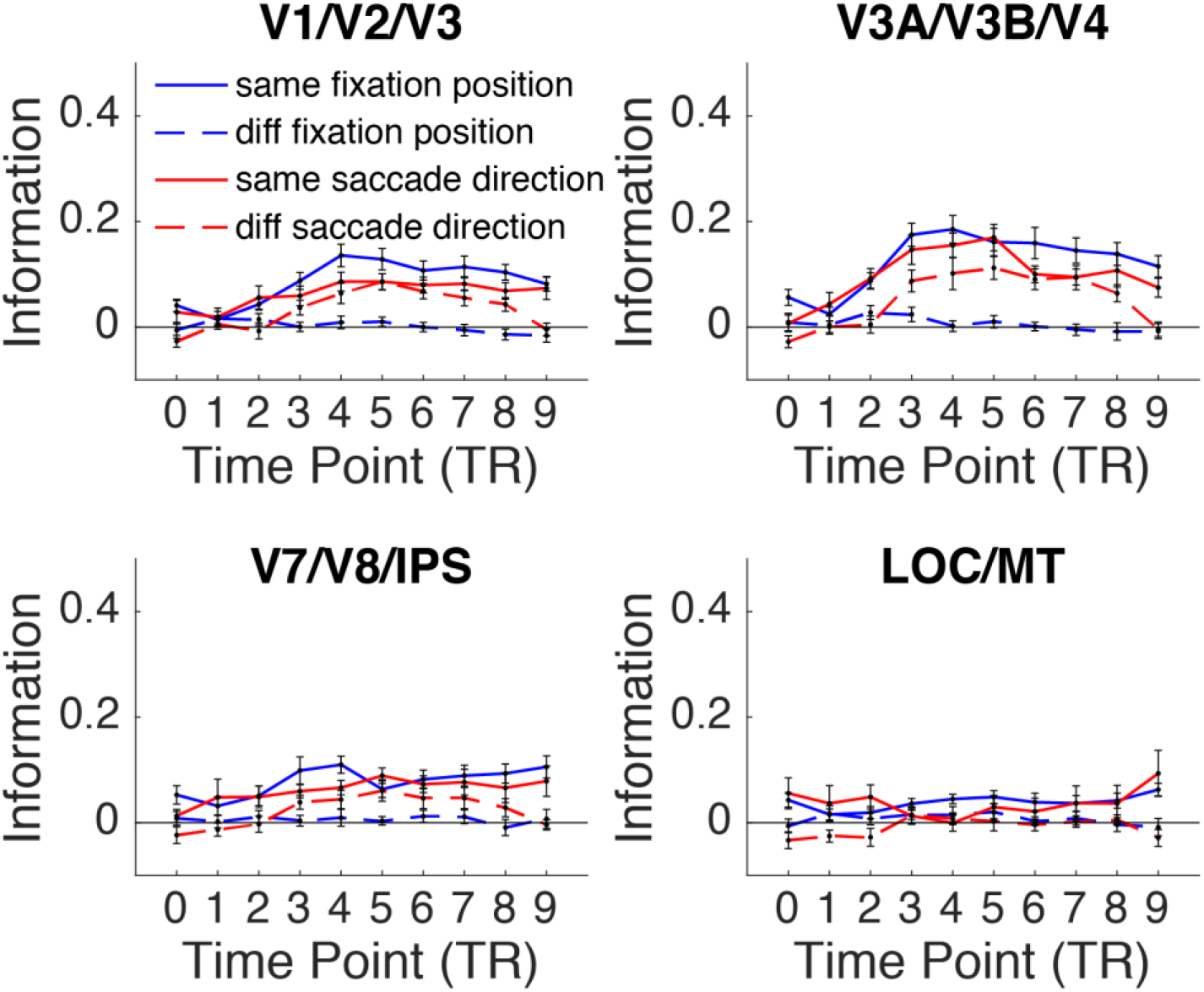
MVPA time course for vertical (Y) location information for each ROI group in the visual hierarchy (depth (z) information timecourses shown in Supplemental Materials). Similar to Figure 4, in most regions location information was dependent on fixation position in no-saccade blocks but tolerant of fixation changes in saccade blocks, even when analyzing timepoint by timepoint. Error bars represent between-subject *SEM*.

## Discussion

In the current study we compared how representations of 3D spatial information in human visual cortex are influenced by eye movements. First, our results during fixation blocks replicated the previous findings of Finlayson et al.(33): MVPA information about the 2D (vertical) location of a stimulus decreased along the visual hierarchy whereas depth location information increased, such that 3D location (depth and 2D) could be decoded from later visual areas. Critically, we revealed that both vertical and depth location could be decoded to a similar extent in dynamic saccade blocks, where participants made a sequence of back and forth saccades while passively viewing the stimulus, as in stable fixation blocks. Follow-up analyses separating fixation positions and saccade directions showed that the representations of 3D spatial locations (both vertical and depth) were *dependent* on fixation position in no-saccade blocks, yet exhibited *tolerance* across fixation changes and different saccade directions in saccade blocks. This pattern persisted even when analyzing timepoint by timepoint, at least for the more robust vertical location information.

### Dependence on fixation position in no-saccade blocks

One of our more striking results was that we barely found any vertical or depth information tolerant of fixation position in the no-saccade blocks. A number of prior studies have reported fixation/eye position effects in neural processing of space. Neurophysiological studies have revealed eye position “gain fields” in parietal cortex – where the neural response to a stimulus depends on the angle of gaze (45). Additionally, information about fixation position has been found alongside retinotopic stimulus location in primate and human visual cortex (25, 46–49), and recent fMRI studies have demonstrated that (static) fixation position can modulate the responses to retinotopic visual stimuli in human visual cortex (47, 50). Our findings are consistent with these studies, revealing a particularly strong modulation by fixation/eye position on the representations of both 2D and depth locations, such that the spatial representations during sustained fixation seemed completely dependent on fixation position, even for spatial dimensions that are not directly affected by the change in eye position (i.e. vertical and depth information, across different horizontal fixation positions).

In the current design, because stimuli always appeared in the horizontal center of the screen, the two different fixation positions by definition resulted in different eye-centered (retinotopic) horizontal locations. Visual regions are known to be organized in retinotopic reference frames (24, 25), and moreover, horizontal information may dominate spatial representations, especially when horizontal location varies across hemifields (33, 51). In the current experiment we could not dissociate the influences of fixation position and eye-centered horizontal stimulus location. However, Finlayson et al.(33) *did* vary horizontal stimulus location alongside vertical and depth location in a very similar design (but with fixation position always held at center). In that study, both vertical and depth location information were found to be at least partially tolerant of differences in horizontal location. This comparison raises the intriguing possibility that representations of vertical and depth locations are dependent on horizontal eye fixation position, but not so much on horizontal stimulus location, even though both manipulations produce the same retinal consequences. However, it is also possible that the combination of fixation position difference and retinotopic / hemifield difference exacerbated the dependence. Regardless, having retinal images “labeled” with the corresponding fixation positions is likely an important mechanism for visual stability (14, 52, 53). Moreover, the strong dependence on fixation position we found in no-saccade blocks is particularly interesting in light of our other finding, that spatial locations *could* be decoded in dynamic saccade blocks, tolerant of changes in fixation position.

### Fixation-tolerant representations of 3D location in saccade blocks

We found that both 2D and depth location information could be decoded in saccade blocks in later visual areas; in fact, we did not observe any degradation of spatial information in saccade blocks compared to no-saccade blocks, even when decoding across blocks with different saccade directions / fixation patterns. We had predicted that horizontal saccades might disrupt depth representations in particular, since horizontal binocular disparity is a main cue for depth perception (indeed, the only cue manipulated in the present experiment), and there is some behavioral evidence showing that horizontal saccades impair depth processing, such as depth judgments of flashed stimuli (42), and memory-guided reaching in depth (43). Yet even though the horizontal saccades may introduce a challenge for preserving accurate stereopsis through binocular disparity, information about stimulus position in depth in human visual cortex was not diminished. If anything, the representations became more robust and tolerant, and this pattern was common across both depth and 2D location.

Of course, the fact that we could reliably decode 3D spatial location from brain activity during the saccade blocks does not mean there was not – or is never -- disruption from saccades. First, our task differed from the behavioral depth tasks in that participants were not required to process the precise depth locations to perform the task; if we had probed more precise depth locations or included an attention to depth component, it is possible there may have been a subtle decrement. Second, in our task there could still have been some transient disruption triggered by the saccade, but the representations could have quickly remapped / recovered after each saccade. Remapping can occur predictively (54) and is at most completed within a few hundred milliseconds after the saccade (5); in the current task the stimulus was present the whole time and saccades were spaced with a 2s gap, so there could have be enough time for the representations of spatial locations to remap and re-stabilize, without the BOLD signal capturing this transient change. Regardless, our findings show that representations of both depth and 2D spatial locations can at least be rapidly updated and preserved across saccades, and – critically – that representations of 3D spatial locations across dynamic saccades are not just a simple aggregation of those representations used during sustained fixations.

### The role of active, dynamic saccades in spatial stability

Our key finding was a pattern of fixation-position-dependency during stable fixation and fixation-position-tolerance during dynamic saccades, for both 2D vertical (y) and depth (z) spatial dimensions. We speculate that the difference between tolerance in saccade versus no-saccade blocks seems to fall on whether active, frequent, and repetitive eye movements were made, such that representations of 3D spatial location (both 2D and depth) *become* more tolerant of fixation position during – and because of – active, dynamic saccades.

As briefly mentioned in the Results section, a potential alternative explanation is one of a temporal blending artifact from the block-design GLM analysis, where the activation pattern on saccade blocks might reflect a combination of left and right fixation patterns, such that the specific saccade pattern/direction becomes meaningless across the block. However, if this were the dominant reason, we would probably expect decoding performance in the saccade blocks to be worse overall compared to no-saccade blocks. That is, given the strong fixation position dependence on no-saccade blocks, if activity patterns were contaminated between left and right fixation periods on saccade trials, we would predict some intermediate cost for decoding. Similarly, the MVPA timecourse analysis argues against this idea. Although our design was not optimized for time-course analyses and individual time points were not independent, the analysis suggests little evidence that the whole-block results could be solely attributable to temporal blending. Instead, the results seem more consistent with tolerance emerging due to some active processing mechanism across saccades; i.e., the blocks in which participants were actively making saccades triggered additional neural signals not present during the sustained fixation blocks, and this could have contributed to greater integration and stability, resulting in more tolerant representations.

The idea of active processing is not new. Self-generated actions are known to influence perception in a variety of ways (55). It has been found that executing or even preparing motor actions can modify perception and representation of 3D space, beyond the direct sensory consequences from the motor actions (see review 44). In the case of eye movements, the motor act generates a corollary discharge, or efferent feedback, signal that feeds back to visual areas and is thought to be a key mechanism for remapping and stabilizing perception (56). This corollary discharge signal can even precede the motor movement itself, allowing for predictive remapping in 2D (57) and 3D (58). Indeed, in a classic study where the eye muscles were paralyzed and eye movements were intended but unable to execute, corollary discharge resulted in false perception of a displacement of the visual scene (59). Similarly, when the thalamus (thought to be critical for relaying efferent feedback) is lesioned, patients may mis-attribute the perceptual consequences of an oculomotor targeting error to external stimulus changes (60). In typical vision, we perceive objects in the world as stationary when we are moving our eyes, but an object displacing the analogous amount on our retinas without an active eye movement would be readily detected as non-stationary. Saccade execution has been shown to contribute to perceptual continuity on the behavioral level (61); consistently, an fMRI study showed that scene representations spanning different views can be integrated across eye movements in scene-selective cortex, but not in the condition where eyes were stable and the scene was moved to mimic the retinal changes induced by eye movements (62) ^2^.

Interestingly, while active corollary discharge mechanisms are thought to play a key role in visual stability, other theories of visual stability focus on more static factors like eye position gain fields, suggesting that spatiotopic (world-centered) neural representations can be formed implicitly by the combination of current retinotopic position and current eye position (14, 47, 53), the latter signal coming from proprioceptive information relaying the position of the eye in the orbit (63). An interesting interpretation of our current findings is that there may be different stability-related signals during active vs static perception, such that 3D location is labeled with eye position information during sustained fixation, but this information is integrated in the context of active eye movements. This maps nicely onto the idea that visual stability across saccades incorporates these two distinct sources of feedback at different timescales: a rapid, predictive corollary discharge signal (triggered by an *active eye movement*), and an oculomotor proprioceptive (*static*) signal that stabilizes more slowly after a saccade (64).

In addition to active remapping via a corollary discharge mechanism, there may be other factors contributing to our observation of 3D location representations becoming tolerant in the context of active, dynamic saccades. These might include factors related to the predictability and expectation of the saccades sequence and/or repetitive retinal changes, as well as questions about whether the increased tolerance could be achieved with a single saccade or is built up over the course of several repeated actions. Follow-up investigation is still needed to further test these possibilities, but all of these considerations indicate the important role of an active observer, in contrast with a passive viewer.

In sum, our findings highlight an important role of active, dynamic saccades on stabilizing 2D and 3D spatial representations in the brain, as well as the possibility that spatial location may be represented differently during active vs static contexts, such that visual cortex may represent the visual world flexibly based on whether sustained fixations or frequent saccades are required.

## Materials and Methods

### Participants

12 right-handed subjects participated in the study and were included in the analyses (6 females, 6 males, mean age 21.42, range 19-34). Three additional subjects were also scanned, but the data were excluded due to excessive amount of head motion, failure to perform the dot-dimming task, and radiologist-detected neuroanatomical abnormality, respectively. All subjects included reported normal or corrected to normal vision, and normal color and depth perception. All gave informed consent and were pre-screened for MRI eligibility. The study protocol was approved by the Ohio State University Biomedical Sciences Institutional Review Board. Before the scanning session, each participant went through a series of pre-screening behavior tasks to assess their depth perception (data not shown; all participants scanned exhibited normal depth perception).

### fMRI acquisition

This study was done at the OSU Center for Cognitive and Behavioral Brain Imaging with a Siemens Prisma 3T MRI scanner using a 32-channel phase array receiver head coil. Functional data were acquired using a T2-weighted gradient-echo sequence with multiband factor of 3 (TR=2000ms, TE=28ms, flip angle 72°). The slice coverage was oriented parallel to the AC-PC plane and was placed to contain full coverage of cerebrum (72 slices, 2×2×2mm voxel). We also collected a high-resolution MPRAGE anatomical scan at 1mm^3^ resolution for each participant. Each participant was scanned in one two-hour session.

### Stimuli and task

The main stimuli and task in the scanner were modified from Finlayson et al.(33), Experiment 1. We used dynamic random dot stimuli (RDS) to stimulate different 3D locations in the participants’ visual field (Figure 1). We used red/green anaglyph glasses with Psychtoolbox’s stereo-mode to achieve depth perception of the RDS stimuli from binocular disparity. The stimuli were small patches (3.3° square) of dynamic RDS located above or below (vertical position) and in front of or behind (depth position) the screen center. The stimulus locations were centered ± 2.7° from the screen center along the vertical dimension, and ± 10 arc min along the depth dimension. A black empty circle (0.06° radius) inside a white dot (0.13° radius) was used as the fixation point. The fixation point could appear in one of two positions: either left or right of the screen center, ± 2.7°, centered vertically and in depth. On half of the blocks (no-saccade blocks), the fixation dot remained in the same position for the entire 16s block. On the other half (saccade blocks), the fixation dot moved back and forth between the two fixation locations every 2 seconds.

To encourage perception of a 3D space, we used a static RDS background field (10.70° square) at the central depth plane of the screen, consisting of light and dark gray dots on a mid-gray background (8 dots/deg^2^, 37% contrast). In addition, we used ground and ceiling lineframes (12.8° × 1.5°) above and below this background RDS, respectively, each spanning ± 13 arc min in front and behind the fixation depth plane. Similar to Finlayson et al. (33), the smaller dynamic RDS stimulus patches comprised black and white dots (100% contrast), with the position of the dots randomly repositioned each frame (60 Hz).

We employed a block-design, with 8 main block types determined by 4 stimulus location conditions (up-front, up-back, down-front, down-back) and 2 fixation/saccade conditions (no-saccade block, saccade block). For ease we abbreviate these 8 main conditions as Fix-up-front, Fix-up-back, Fix-down-front, Fix-down-back, Sac-up-front, Sac-up-back, Sac-down-front, Sac-down-back, where Fix-up-front means a no-saccade (sustained fixation) block where the stimulus appeared in the upper (vertical) and front (depth) position. Each run included 16 stimulus blocks, two blocks per main condition, to maintain the same power as in Finlayson et al.(33). We arranged the order of block conditions in a pseudo-random fashion (i.e., balanced Latin square design) for each run. We added three null-stimulation baseline blocks in each run, where there were no RDS stimuli presented, just a fixation dot. The null blocks were added at the beginning (Block 1), middle (Block 10), and end (Block 19) of each run. All blocks lasted 16s, with a 2s inter-block interval, and each run lasted 360s. Each subject completed 8 runs of the task.

In all conditions, participants were instructed to fixate on the fixation dot and perform a dot dimming task at fixation, detecting when the black empty circle was filled in to be a solid black circle. When the fixation changed its location, participants were instructed to move their eyes to follow it. There were two possible fixation positions (left and right). The fixation position for each no-saccade block, and the initial fixation position for each saccade block, were selected as follows: In each run, the sequence of the fixation positions was determined so that there were no inter-block eye movements (i.e., the final fixation position of the previous block was the same as the initial fixation position of the current block). This was important to ensure that saccades were only executed during saccade blocks. The sequence was further constrained to ensure that both left and right fixation positions were equally likely across the 8 main conditions over the course of the experiment. The no-saccade conditions were balanced such that for each of the four no-saccade stimulus location conditions, there was one block per run with left fixation position and one block per run with right fixation position. Because each saccade block involved a repetitive sequence of 8 alternating saccades, an equal number of leftward and rightward saccades were always included in each saccade block, though the saccade direction was dictated by the initial fixation. For saccade blocks, the distribution of initial fixation positions could not be fully counterbalanced within each run, given the constraint above. E.g., within a given run, the two blocks of the Sac-up-front condition might have both been assigned a left initial fixation position; but across the total 8 runs of the experiment, an equal number of left and right initial fixation positions were assigned to each condition.

Participants wore red/green anaglyph glasses to view the 3D stimuli, and were instructed to flip the direction of the glasses midway through the experiment (after four of eight runs), in order to control for low-level stimulus differences in the MVPA analysis due to the color presented to each eye, per Finlayson et al.(33). When the glasses were flipped, the stimulus code adapted to reflect the current glasses direction and stimulate the correct depth location (i.e., front or back).

All stimuli were generated with Psychtoolbox (65) in Matlab (MathWorks). Stimuli were displayed with a 3-chips DLP projector onto a screen in the rear of the scanner (resolution 1280×1024 at 60Hz). Participants viewed from a distance of 74cm via a mirror above attached to the head coil.

### Eye tracking

Eye gaze positions were recorded inside the scanner throughout the experiment, using an MRI-compatible Eyelink remote eye-tracker at 500 Hz. Eye position data were used to ensure the participants kept their eyes on the fixation point and made eye movements following the fixation change. Due to the red-green anaglyph glasses interfering reflection, the eye-tracking calibration was not always reliable for all subjects. In circumstances where reliable eye position data were not able to be recorded, the experimenters could observe the subject’s eye through the camera video and/or use the behavior performance of the dot dimming task to ensure that the participants were making eye movements as intended.

### fMRI preprocessing and analyses

The fMRI data were preprocessed with Brain Voyager QX (Brain Innovation). All functional data were corrected for slice acquisition time and head motion and temporally filtered. Runs with abrupt motion greater than 1mm were discarded from later analyses. Spatial smoothing of 4mm FWHM was performed on the preprocessed data for univariate analyses, but not for multivariate (MVPA) analyses. Data of each participant were normalized into Talairach space (66). We used FreeSurfer to segment the white matter/gray matter boundaries for each participant’s anatomical scan, to inflate and flatten each hemisphere into cortical surface space.

A whole-brain random-effects general linear model (GLM), using a canonical hemodynamic response function, was used to calculate beta weights for each voxel, for each condition and participant. More details are given in the multivariate pattern analysis (MVPA) section. All GLM data were exported to Matlab using Brain Voyager’s BVQXtools Matlab toolbox, and all subsequent analyses were done using custom code in Matlab.

### Functional localizers and Regions of Interest (ROIs)

In addition to the main task, each participant also completed 3 runs of functional localizers and 4 runs of 2D retinotopic mapping. We identified a priori ROIs similar to Finlayson et al.(33):

We defined retinotopic areas V1, V2, V3, V3A, V3B, V4, V7, and V8 using a standard retinotopic mapping paradigm with a rotating wedge with high-contrast radial checkerboard patterns (22). The 60° wedge stimulus covered eccentricity from 1.6° to 16° and flickered at 4 Hz. It was rotated for 7 cycles with a period of 24s per cycle in each run, either clockwise or counterclockwise. Participants’ task was to fixate at the center fixation on the screen and press the button when the fixation dot changed color. After preprocessing, the brain data of retinotopic mapping runs were analyzed in custom Matlab code and were projected onto the flattened cortical surface maps in Brain Voyager, and boundaries between the retinotopic areas were delineated.

For each individual participant, we also defined the object-selective LOC (2 localizer runs) and motion-sensitive MT+ (1 localizer run). The LOC localizer task included blocks of grayscale real-world objects and scrambled objects (12°×12°) presented at the center of the screen. Participants performed a one-back repetition task, where they pressed a button whenever the exact same stimulus image was presented twice in a row. The object-selective LOC region was defined with an object > scrambled contrast. For the MT+ localizer task, participants fixated at the center of the screen and passively viewed blocks of either stationary or moving random dot displays. The stimuli were full screen dot patterns, and the moving patterns alternated between concentric motion towards and away from fixation at 7.5 Hz. The motion-sensitive MT+ area was defined with a moving > stationary contrast. We additionally localized a visually sensitive intraparietal sulcus (IPS) ROI using data from the LOC localizer task (All > Fixation contrast) in conjunction with anatomical landmarks.

To compare with findings in Finlayson et al.(33), we grouped the ROIs in a similar way, according to their relative positions along the visual processing hierarchy: early visual areas V1, V2, and V3; intermediate visual areas V3A, V3B, and V4; later visual areas V7, V8, and IPS; and category selective areas LOC and MT+. For our main analyses, MVPA information was calculated for individual ROIs and subjects and then averaged across the ROIs in each group. Analyses with all individual ROIs are shown in the supplemental materials.

### Multivoxel pattern analyses (MVPA)

Multivoxel pattern analyses (MVPA) following the split-half correlation method (67) were performed for ROI-based as well as whole-brain searchlight analyses. (Searchlight results and methods are presented in the supplemental materials.) Our main approach of quantifying vertical and depth location information was similar to Finlayson et al.(33).

For our primary analyses, we compared vertical and depth information in the no-saccade versus saccade blocks. We conducted MVPA on the 8 main conditions (4 stimulus location conditions × 2 fixation/saccade conditions). We split the data into two halves based on the direction of the anaglyph glasses, correlating multivoxel patterns of activity for each of the 8 main conditions in the first 4 runs (RG runs, red color over the left eye) with each of the 8 main conditions in the last 4 runs (GR runs, green color over the left eye). We ran GLMs with the 8 main conditions as our 8 regressors of interest for each split-half dataset. The beta weights for each voxel were normalized (within each dataset) by subtracting the mean response across all conditions for that voxel from the responses to individual conditions. Next, the voxel-wise response patterns for each of the 8 conditions in the RG runs were correlated with each of the 8 conditions in the GR runs, generating an 8-by-8 correlation matrix (Figure 2A). The correlations were converted to z-scores using Fisher’s r-to-z transform.

We then quantified the amount of vertical (Y) and depth (Z) location information contained within each ROI for each subject, as follows. First, the cells in the correlation matrix were characterized according to whether they reflected the same or different Y location, same or different Z location, and same or different fixation/saccade condition (no-saccade or saccade). For example, the Sac-up-back (RG-runs) × Sac-down-back (GR-runs) correlation would be characterized as same fixation/saccade condition, different Y, same Z (1 0 1). Then, for each type of information, we averaged across all of the “same” cells for that type of information, and all of the “different” cells (Figure 2A), and the “same” minus “different” correlation difference was taken as a measure of the amount of “information” about that property. E.g., y-information was quantified as the difference in correlation between all conditions that shared the same Y position (- 1 -) versus differed in Y position (- 0 -). This standard approach (67) is based on the rationale that if an ROI contains information about a certain type of location, then the voxel-wise response pattern should be more similar for two conditions that share the same location than differ in location. Y and Z information were calculated in four ways: for all blocks (ignoring saccade information; Figure 2), within no-saccade blocks only (Figure 3A), within saccade blocks only (Figure 3B), and cross-decoded *between* no-saccade and saccade blocks (Figure 3C). All analyses were performed within each ROI and subject, and then averaged into the ROI groups where applicable. Standard frequentist within-subject statistics were performed comparing whether the different types of information were significantly different from zero and/or each other.

As a second stage of analysis, we broke down the conditions further to compare within-versus across-fixation location information in no-saccade blocks, and within-versus across-saccade-direction location information in saccade blocks (Figure 4). For each of these analyses we used different GLMs in order to maximize power for the comparisons of interest. The main reason for different GLMs is that no-saccade fixation locations were perfectly balanced within each run, but saccade direction was only balanced across runs, as described earlier.

For the no-saccade breakdown (Figure 4A), we analyzed left and right fixation position blocks as separate conditions, running the split-half GLMs with 12 conditions: The 4 original no-saccade conditions were now broken down into 8 conditions, based on whether the fixation position was on the left or right for that block. These 8 no-saccade conditions were coded as FixLeft-up-front, FixLeft-up-back, FixLeft-down-front, FixLeft-down-back, FixRight-up-front, FixRight-up-back, FixRight-down-front, FixRight-down-back. The original 4 saccade-block conditions (Sac-up-front, Sac-up-back, Sac-down-front, Sac-down-back) were modeled in the GLM, but ignored in the MVPA analysis. We used the same approach described earlier to generate split-half correlation matrices, here between the beta weights of the 8 no-saccade conditions in the RG-runs and the 8 no-saccade conditions in the GR-runs, for each subject and ROI. Y and Z location information were each then calculated for two contexts, using different subsets of the correlation matrix: (1) across no-saccade blocks with the same fixation position (“no-saccade same fixation”), and (2) across no-saccade blocks with different fixation positions (“no-saccade different fixations”).

For the saccade direction breakdown (Figure 4B), we performed an analogous analysis with 12 conditions, here separating the saccade blocks based on initial fixation position, such that the 4 original saccade conditions were now broken down into 8 conditions, coded as SacLeft-up-front, SacLeft-up-back, SacLeft-down-front, SacLeft-down-back, SacRight-up-front, SacRight-up-back, SacRight-down-front, SacRight-down-back. The original 4 no-saccade conditions (Fix-up-front, Fix-up-back, Fix-down-front, Fix-down-back) were modeled in the GLM but ignored in the MVPA analysis.

Similar to above, we calculated Y and Z information across (1) saccade blocks with the same saccade direction (“saccade same direction”) and (2) saccade blocks with different saccade directions (“saccade different directions”). However, because of the difficulty noted earlier in balancing initial saccade directions within runs, we could not use the simple RG and GR split-half approach, so we adopted an iterative random sampling split-half approach, similar to Lescroart et al., 2016 (68). In this approach, we ran GLMs for each run separately, each containing only 4 of the 8 new saccade-breakdown conditions (for each stimulus location condition, only one of the initial saccade directions was present per run). Then, for each of the 8 saccade-breakdown conditions, we randomly split all the runs containing this condition into two halves in each iteration and averaged the beta weights for that condition across the runs included in each half. This gave us two split-half voxel-wise response patterns for each of the 8 conditions (per subject and ROI), which were used analogously to the RG and GR split-half patterns, to create correlation matrices for the MVPA analyses. We performed 100 iterations of the random splitting procedure, and the information results for these 100 iterations were averaged for each subject.

### Multivariate pattern time-course analyses

In order to explore the dynamic representations of spatial locations in both no-saccade and saccade blocks, we performed MVPA time-course analyses using finite impulse response (FIR) GLMs with 10 timepoints; one timepoint per 2 sec TR, where timepoint zero (TP0) corresponds to the start of each block (i.e., the onset of the random dot stimulus). To be able to directly compare the time-courses of no-saccade conditions (same-fixation vs different-fixations) and saccade conditions (same-saccade-direction vs different-saccade-direction) in the same analysis, we included all 16 breakdown conditions (4 stimulus locations × 2 no-saccade fixation locations + 4 stimulus locations × 2 saccade direction conditions) as regressors of interest, using the iterative random split-half procedure described above. For each timepoint, we calculated Y and Z information for each tolerance context, as well as fixation position information for no-saccade and saccade blocks, for each ROI and subject. Note that this analysis is substantially less powered than our main analyses, and as a result may only detect larger effects; it is presented primarily as an exploratory analysis to complement the main analyses.

## Acknowledgements

This study was funded by National Institutes of Health grant R01-EY025648 (JG). The authors thank Andrew B. Leber for suggestions on the analyses, and members of the Golomb Lab for helpful feedback.

## Supplemental Materials

### Table of contents

Figure S1. Whole-brain searchlight analyses of vertical and depth location information.

Figure S2. Y and z location information in each individual ROIs along the dorsal and ventral pathway in left and right hemispheres.

Figure S3. MVPA decoding in the time course for information about (initial) fixation position, vertical location, and depth location.

Figure S4. Univariate analyses of individual ROIs for each stimulus location condition.

**Figure S6.**
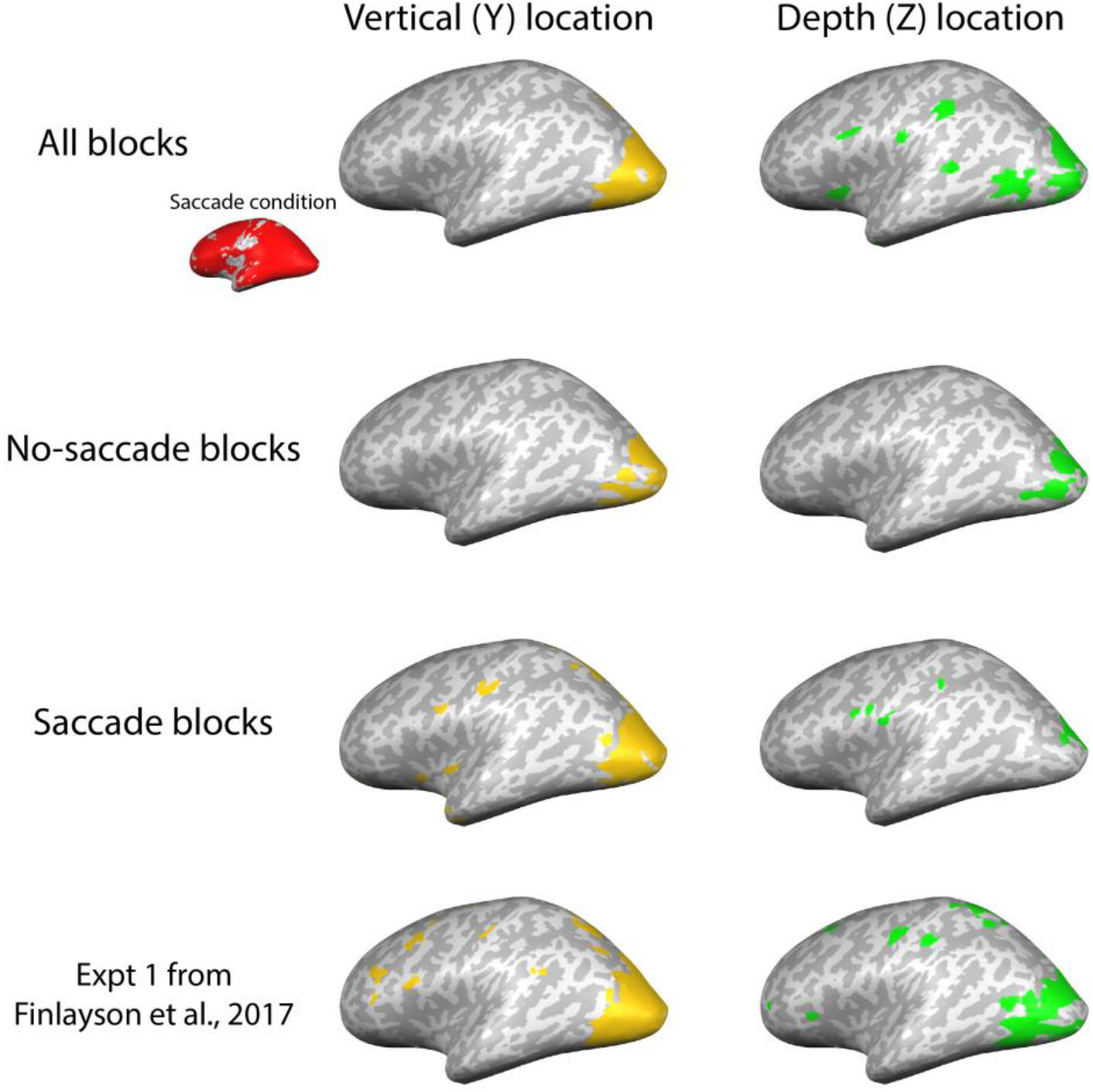
Searchlight results. Inflated brains show voxels with significant vertical (y) and depth (z) location information, corresponding to the comparisons in main text Figures 2–3. Top row: collapsing across all blocks; 2^nd^ row: for no-saccade blocks; 3^rd^ row: for saccade blocks; 4^th^ row: data re-analyzed from Experiment 1 of Finlayson et al., 2017 and plotted in the same manner for comparison. The top row inset also shows voxels with significant information about no-saccade vs saccade condition in red. The searchlights are shown mainly for illustrative purposes; we probe these patterns more quantitatively with the a priori ROI analyses in the main text. It should be noted that although we applied similar analysis procedures for exploratory searchlight and a priori ROI analyses, some divergence may be expected due to the fact that the searchlight analyses averaged across subjects in a common space and thus only picked up voxels consistent across subjects, whereas the functional ROIs were localized and defined based on each individual’s functional anatomy. Methods: For each participant, we iteratively searched through the brain conducting MVPA within a “moving” ROI defined as a sphere of radius 3 mm (~100 voxels). On each iteration, the ROI was chosen as a sphere centered on a new voxel, and multivoxel correlation analyses were performed in the same way as described in the main text. The magnitudes of Y and Z information (as defined by the z-transformed “same” minus “different” correlation differences) were then plotted for each voxel, creating a z-map for each type of information for each participant. These participant-level maps were then spatially smoothed with a 3 mm FWHM kernel. To generate the group-level maps shown in the figure, we used one-sample t-tests to identify clusters containing significant information about each type of information. The resulting t-maps were thresholded at p < .05 and cluster size > 25 contiguous voxels; voxels are color-coded according to whether or not they were significant for each type of information.

**Figure S7.**
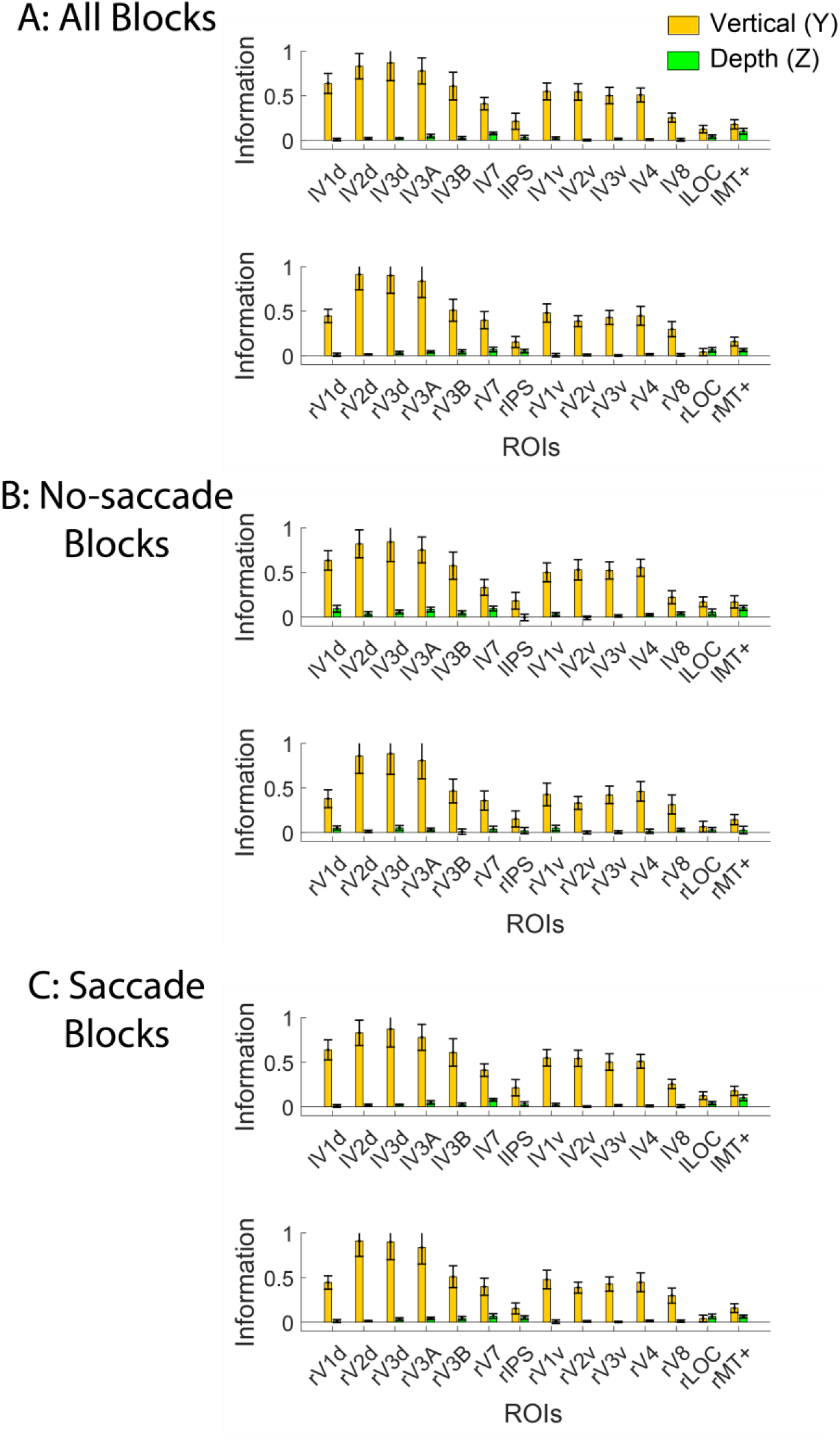
3D location information in individual ROIs. MVPA analyses are shown for vertical (y) and depth (z) information in all blocks (A; similar to Figure 2), no-saccade blocks (B; similar to Figure 3A), and saccade blocks (C; similar to figure 3B). ROIs are named based on the hemisphere (l/r) and dorsal/ventral division when appropriate: e.g., lV1d is area V1 in the left hemisphere, dorsal stream. Error bars represent between-subject SEM.

**Figure S8.**
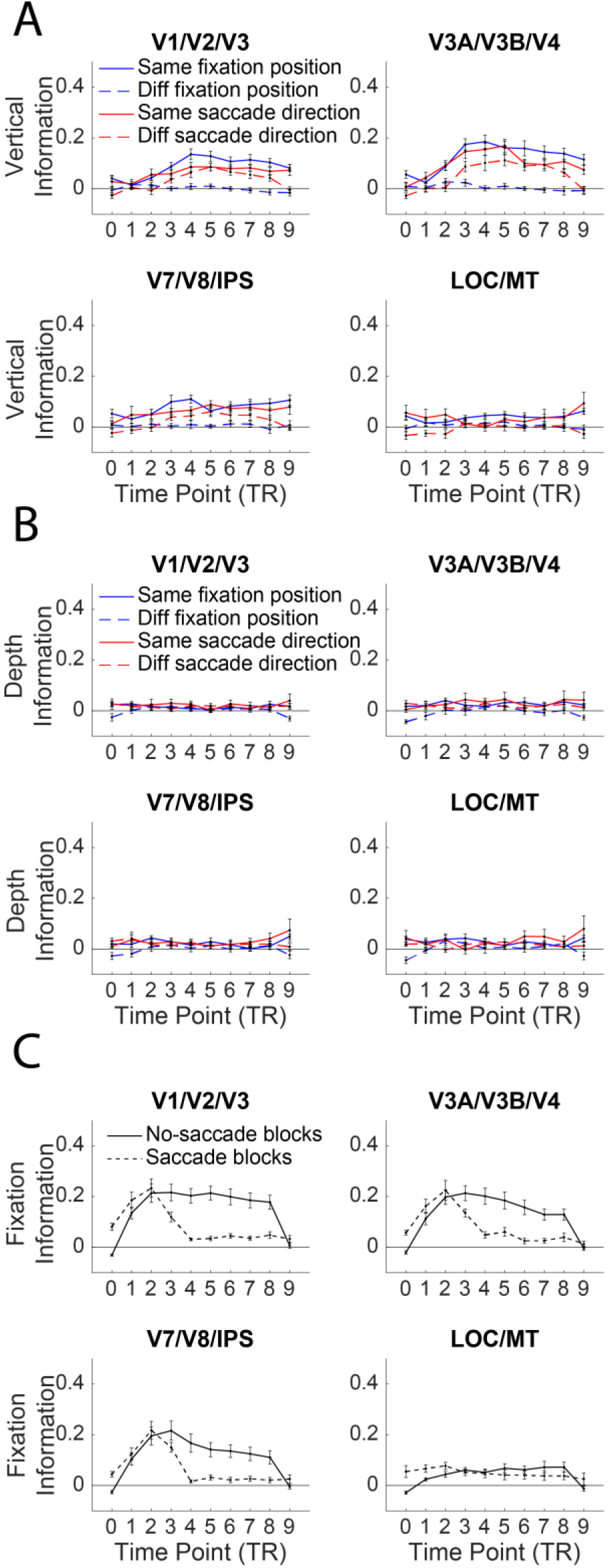
Full MVPA time course results. (A) MVPA timecourse for vertical (y) location information for each ROI group in the visual hierarchy, replotted from main text Figure 5. (B) Same as A, but for depth (z) location information. (C) MVPA timecourse for information about fixation position (left vs right) in no-saccade blocks (solid line) and about initial fixation position in saccade blocks (dashed line). Error bars represent between-subject SEM.

**Figure S9.**
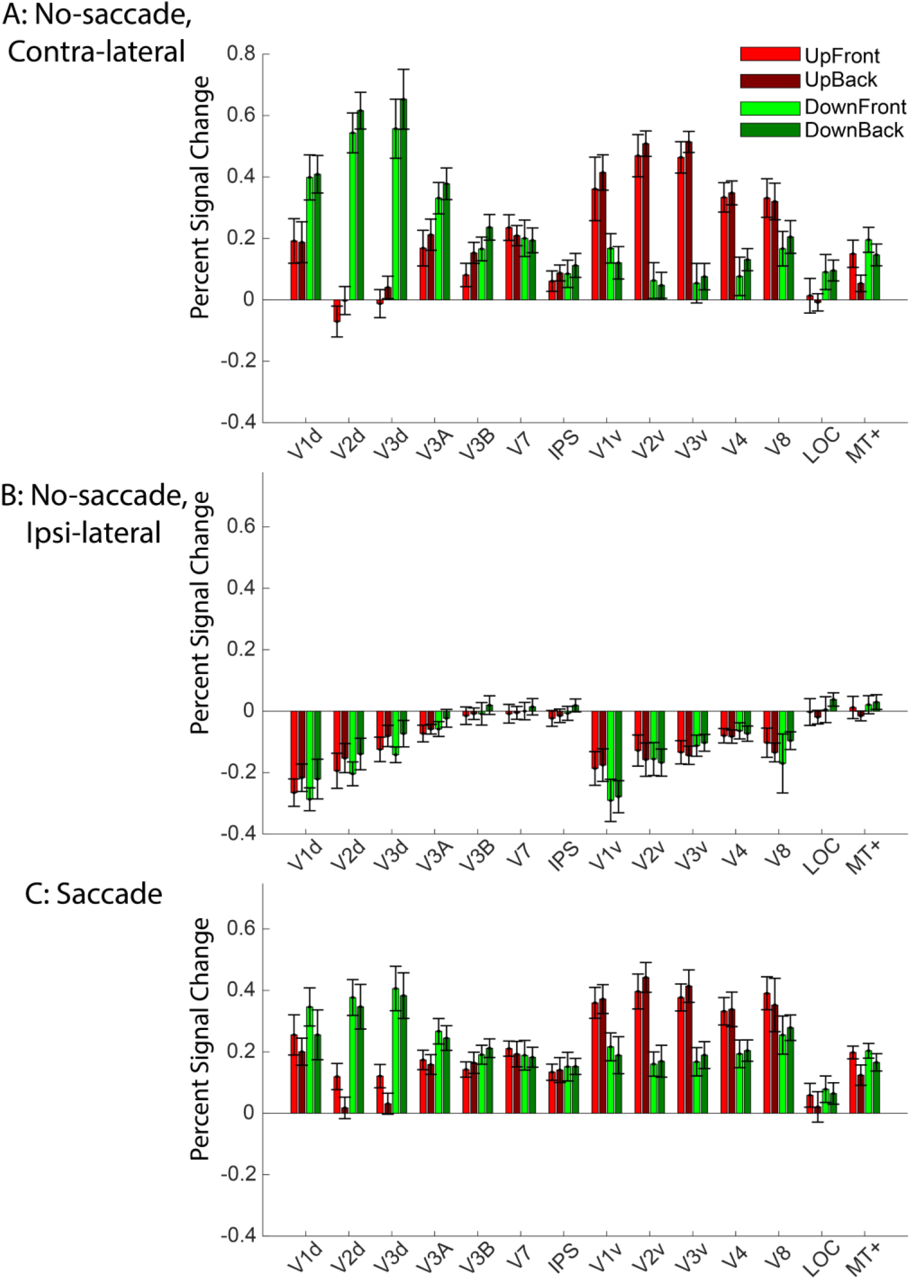
Univariate results. BOLD response (percent signal change) evoked by each of the 4 3D stimulus locations (vertical x depth) for individual ROIs. A and B: No-saccade blocks, plotted separately for stimuli appearing in contralateral and ipsilateral visual fields. For each ROI, we first separated by hemisphere and separately plotted the data for left and right fixation positions, then re-coded them as ipsi-lateral or contra-lateral so we could average across left and right hemispheres for each ROI. The univariate data confirm strong contra-lateral preferences, along with expected vertical preferences for dorsal and ventral ROIs. There were no significant depth differences between the univariate responses to front vs back stimulus locations. C: Univariate responses plotted for saccade blocks (averaged across saccade direction). Error bars represent between-subject SEM.

## Footnotes

1 Y information was significant in all ROIs in both same-direction and different-direction conditions *t*’s≥2.229, *p*’s≤.048, Cohen’s *d*’s≥0.644, except for different-direction category-selective regions, *t*_11_=1.687, *p*=.120, Cohen’s *d* =0.487. Z information was significant in all ROI groups in the different-direction condition, *t*’s≥2.525, *p*’s≤.028, Cohen’s *d*’s≥0.729; however, z information in the same-direction condition was not significantly above zero in any of these regions, *t*’s≤2.121, *p*’s≥.057, Cohen’s *d*’s≤0.612. (The difference of z information between same-direction and different-direction conditions was significant in early and later visual ROIs, *t*’s≥2.814, *p*’s≤.017, Cohen’s *d*’s≥0.812, but not in intermediate and category-selective ROIs, *t*’s≤0.453 *p*’s≥.660, Cohen’s *d*’s≤0.131.)

2 In the current study we did not include a comparison condition where the fixation was kept stable and the stimulus jumped between left and right sides every 2s. Although on the surface this would seem like a compelling way to differentiate whether location representations become more stable due to dynamic saccades specifically, versus other dynamic retinal changes, it would have been a problematic comparison because as noted earlier, Finlayson et al.(33) already found that vertical and depth location information were partially tolerant to horizontal location when the eyes remain fixated, even in the absence of dynamic context.

